# The converse effects of speed and gravity on the energetics of walking and running

**DOI:** 10.1101/201319

**Authors:** S.J. Hasaneini, R.T. Schroeder, J.E.A. Bertram, A. Ruina

## Abstract

The energetic cost of transport for walking is highly sensitive to speed but relatively insensitive to changes in gravity level. Conversely, the cost of transport for running is highly sensitive to gravity level but not much to speed. Gait optimization with a minimally constrained bipedal model predicts a similar differential energetic response for walking and running even though the same model parameters and cost function are used for both gaits. This challenges previous assertions that the converse energetic responses are due to fundamentally different energy saving mechanisms in each gait. Our results suggest that energetics of both gaits are highly influenced by dissipative losses occurring as leg forces abruptly alter the center of mass path. The observed difference in energetic consequence of the performance condition in each gait is due to the effect the movement strategy of each gait has on the dissipative loss. The optimization model predictions are tested directly by measuring metabolic cost of human subjects walking and running at different speeds in normal and reduced gravity using a novel reduced gravity simulation apparatus. The optimization model also predicts other, sometimes subtle, aspects of gait such as step length changes. This is also directly tested in order to assess the fidelity of the model’s more nuanced predictions.

## Introduction

The pattern of movement in a gait is determined by the central nervous system (CNS), which selects a specific coordination strategy from a vast array of possible alternatives, some more viable than others. It would be useful to understand the factors that influence the selection of the gait coordination strategy and the specific gait parameters involved (such as step length and frequency for a given speed) as this would provide substantial insight into predicting behavior and defining limits in all functional environments encountered – be they moving on other planets or contending with the restrictions imposed by physical disabilities on Earth.

The current prevailing consensus suggests that preferred gait coordination patterns exist to facilitate exchange between energy forms so that energy can be recovered from within the stride for use in another portion of the cycle (Biewener, 2006, Cavagna et al., 1977, Saibene, 1990). Although such exchanges can be observed, evidence is accumulating that energy recovery *per se* may not be as important a determinant of gait coordination strategy as has previously been assumed (Kuo, 2001, 2002, Ruina et al., 2005, Bertram and Hasaneini, 2013).

If exchange between energy forms is not a critical determinant of the selected coordination pattern in gait, what does influence the choice of movement strategy in any given circumstance? One alternative suggests that gait solutions are optimized compromises between competing physical penalties (Kuo, 2001, Hasaneini et al., 2015, McGeer, 1990a, b). Some insight into the factors influencing the CNS strategy can be inferred by using predictive models as quantitative hypotheses, where model predictions of gait coordination in specific circumstances can be tested against the movement patterns selected by human subjects negotiating the same physical conditions. By allowing the model to independently ‘discover’ the optimal coordination pattern under minimal constraints, the influence of the optimization function can be quantitatively evaluated. A clearly specified model makes the reason behind the specific optimization solution accessible to analysis. This is particularly insightful for conditions that differ from those normally encountered. A model that responds to unusual operational circumstances in a manner similar to that of the human system may well point to the factors important in explaining why the CNS selects the coordination strategy it does.

Observation of normal walking and running makes human locomotion appear stereotyped (Pribe et al., 1977), suggesting a pre-programmed control pattern. However, when faced with even modestly unusual challenges, alternate solutions are routinely and consistently selected; for instance, when step frequency or step length is artificially constrained rather than speed, different speed-frequency relationships are utilized for both walking and running (Bertram et al., 2001, Gutmann et al., 2006). It is not yet clear why the locomotion control system adapts to unusual circumstances as it does, but a number of studies point toward the important influence of energetic cost optimization (Bertram, 2005, Selinger et al., 2015). One ‘unusual’ circumstance in which locomotory control obviously adapts, albeit in a manner that has defied easy explanation, is that of reduced gravity. For decades it has been recognized that many aspects of walking and running change in substantially different ways as a consequence of moving within either an actual (Cavagna et al., 2000, De Witt et al., 2010) or a simulated reduced gravity environment (Farley and McMahon, 1992, Newman et al., 1994, Donelan and Kram, 2000). For example, two and a half decades ago, Farley and McMahon (1992) demonstrated that under simulated partial^1^ reduced gravity conditions the metabolic cost of transport (COT, energy expenditure per unit body mass and unit distance travelled) decreases in running more dramatically than it does for walking (Fig. 1). In normal gravity, walking at 1 m/s is more energetically cost-effective than running at 3 m/s (typical preferred walking and running speeds). However, due to the greater influence of gravity reduction on the energetic cost of running, the scenario is reversed at low gravity levels (e.g. on the Moon), and running becomes more energetically favorable.

**Figure 1:**
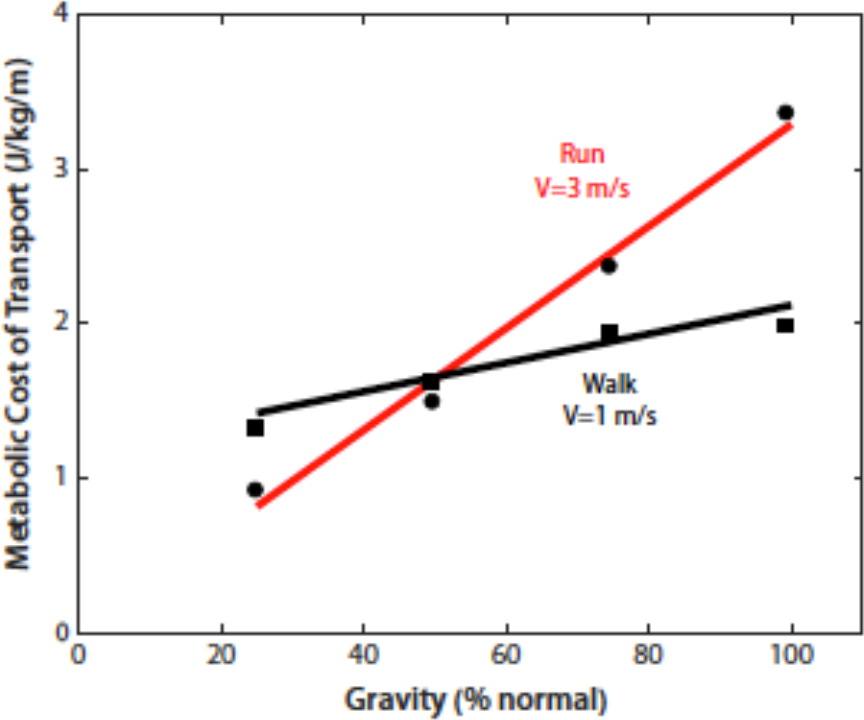
Metabolic cost of transport (net metabolic energy expenditure per unit body mass and unit distance traveled) for walking and running at a range of simulated reduced gravity levels (reproduced from Farley and McMahon, 1992). Energetic cost decreases for both gaits as gravity is reduced, but the effect is more dramatic in running that in walking. As our common experience would indicate, running demands more energetic cost than walking in normal (Earth) gravity. However, due to the different cost reduction rates of these gaits, for gravity below approximately 50% of Earth’s running becomes less energetically costly than walking to travel a given distance.

The differential response of walking and running cost to gravity reduction may appear unexpected, but is simply another aspect of the inherent differences existing between the movement strategies referred to as ‘walking’ and ‘running’. For instance, it has long been recognized that the COT of walking is highly sensitive to changes in speed whereas that of running is remarkably insensitive, at least over a broad range of normal running speeds (Margaria et al., 1963, Wong and Donelan, 2017). Although speed and gravity have different effects on these two gaits, there is currently no conclusive explanation for why walking COT should be sensitive to speed and relatively insensitive to gravity change while running COT is relatively insensitive to speed changes but highly sensitive to gravity reduction. We see these differences as clues to the underlying dynamics of the two gaits and suggest that a model explaining these differential responses will provide a comprehensive understanding for why these movement strategies are used as they are and have the energetic requirements/advantages observed.

Whereas many walking and running motor control models emphasize the driving signals that generate the observed motions, our approach in determining key control mechanisms and their consequences differs fundamentally. We treat gait coordination as a constrained optimization problem and recognize that the integration of the system with its physical environment plays an important role in providing dynamic optimization opportunities. From this, we predict that the motor control system will spontaneously select a coordination pattern that, for the given objective, optimizes (or at least tends toward optimizing) the dynamic interaction of the whole system with the substrate on which it moves.

Working from this approach, we have used gait optimization with a simple, minimally constrained bipedal model based on limb proportions and mass distributions of an average human. Utilizing minimal constraint allows the optimal response to freely emerge without being determined by imposed motion expectations, guidance or performance constraints (other than the optimization criteria). However, successful prediction of the movement strategy and energetic consequences in human subjects operating over a range of speeds and in both normal and unusual circumstances would suggest that the determinants of the model and the human system have a similar basis. Identifying even some of the key determinants of the motor control response in this way would provide substantial insight into how and why human motor control responds as it does during locomotion.

A variety of system features can be optimized, of course, and it is likely that many have at least some effect on the final coordination pattern. However, when considering the circumstances of standard locomotion, where the objective is to move from one location to another at a given speed (without undue effects of instability), we expect that the COT will be a critical determinant of the most effective movement strategy. This is based on the old and intuitive perspective that acting with minimal work is a perpetual law of Nature (Borelli, 1681, Alexander, 2001, Ruina et al., 2005). Optimization models have been providing substantial insight into the fundamentals of gait selection (Srinivasan and Ruina, 2006). Extending the Srinivasan and Ruina (2006) model our previous work (Hasaneini et al., 2013) used a COT calculated from a work-based energetic cost and predicted some naturally selected features of human gaits in normal functional circumstances, including the optimality of walking at slow speeds and running at fast speeds, presence of a preemptive push-off at the step-to-step transition between support legs in both level-ground and uphill walking (and its absence in downhill walking), landing on a near-vertical leg in running, step-length trends with speed, application of swing-leg retraction in both walking and running, and appropriate torso leaning in uphill or downhill gaits.

In the current study we use this model to predict how an individual should respond to both speed and reduced gravity conditions in order to optimize COT. We then compare these predictions to the response of human subjects walking and running at different speeds and in different simulated reduced gravity environments. We evaluate the predictive capacity of the model using two studies, both of which employ a novel reduced-gravity simulation apparatus. First, we extend the data of Farley and McMahon (1992) by evaluating the energetics of walking and running over a range of speeds in normal and simulated reduced gravity. Second, we rigorously analyze the relationship between step length and gravity reduction for both walking and running. If the model’s predictions match observations of human gait, it is likely that the optimal gaits of the model and human gait coordination are influenced by similar determinant factors, especially considering the success of previous predictions in normal gravity (Hasaneini et al., 2013). Since the model is relatively simple, it is possible to probe the model *post hoc* to determine the physical advantages behind the movement patterns selected (or the disadvantages of utilizing alternate movement patterns). This provides a means of understanding the motivations behind the movement adaptations selected and identifying why these have the energetic consequences they do.

## Methods

### The model

The model, shown in Fig. 2, has been described in detail previously (Hasaneini et al., 2013). Briefly, it includes a torso, flat feet, and telescoping legs equipped with rotational hip and ankle joints. The mass distribution and body-segment dimensions are those estimated for a human subject with 75 kg body mass and 176 cm height (Winter, 2005). All joints (linear and rotational) are powered. The stance-leg telescoping actuator can only apply extensional forces. Each hip-motor applies torque between the torso and the corresponding leg.

**Figure 2:**
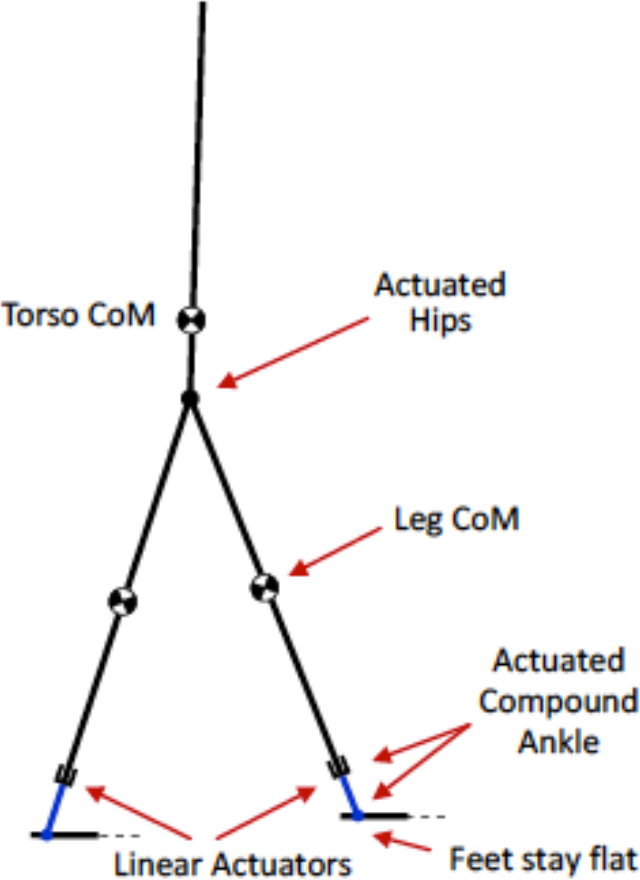
The biped model used for optimization analysis. The torso and legs have distributed mass. All body parts are rigid, and there is no energy recoil (elasticity) in the model. Feet are massless and always stay flat. The ankle is a compound joint, consisting of a (i) rotary joint between the foot and the leg and (ii) a prismatic joint along the leg. A telescoping actuator along each leg applies extensional force along the stance leg, and a motor between the torso and each leg applies torque at each hip joint.

The flexing knees of the human leg are not included in the model. However, leg-length extension and contraction, which are partly due to extension and flexion of the knee in humans (Lee and Farley, 1998), are functionally represented by the telescoping leg. As well, the model allows for evaluation of the ankle’s role during plantar flexion of the foot. Although torque production is allowed for at the ankle axis (as a lever function of the foot), the emerging energy optimal gaits never utilize this mode of actuation. Instead, all optimizations of the model selected leg-length changes (employing the linear actuator) as opposed to active rotation at the ankle. Consequently, the model can be reduced to a point-foot biped without any expectation of a different optimal behavior. This suggests that much of human ankle action during gait is to provide push-off and contribute to appropriate leg length changes over the course of the stride (Hasaneini et al., 2013).

Our model is slightly more complex than the minimal biped studied by Srinivasan and Ruina (2006). Their minimal model had massless legs and, consequently, the energetic cost always converged on a zero or lowest-allowed step length (regardless of speed and gravity level) since there is no energetic penalty for leg swing. One way around this unrealistic solution is to force a step length constraint. However, in the current case we would like to study the effect of gravity on factors contributing to realistic energetic cost and details of stride parameter choices, such as preferred step length. Thus, it is inappropriate to predefine step length. Therefore, we use legs with realistic mass. As a result, energy consequences due to inertia influence the optimization and the model outputs a more realistic energetic cost as well as optimal stride parameters, such as step length. Also, unlike the minimal biped model, our model has a torso, allowing for a separate actuator between each leg and the torso, similar to the human body.

An implementation of trajectory optimization in MATLAB using SNOPT (Gill et al., 2005) finds the optimal gaits for which the COT is minimized, subject to a given gait speed and gravity level. Two types of periodic gait are considered: walking and running. The type of gait is predefined for the optimization, only by defining the sequence of events and phases: heel-strike, double-stance, toe-off, single-stance, and again heel-strike on the opposite foot for walking, and touch-down, single-stance, takeoff, flight, and again touch-down on the other foot for running. No *a priori* assumptions are made regarding kinetic and kinematic parameters such as the joint angles, duration of double-support (extended or instantaneous), type of foot-ground contact (collisional or collision-less), step length, step period, *etc*.

The COT (the objective function) is calculated from COT=E_step_/(m L_step_), where m is the total body mass, L_step_ is the step length chosen by the optimization, and E_step_ is the energetic cost of each step. E_step_ is calculated using a work-based cost model in which the energy expended by each actuator is scaled for positive and negative mechanical work via work-efficiency constants. If W^+^ and W^−^ are the net positive and negative work performed by all actuators in one step, and e_1_ and e_2_ are the corresponding work efficiency constants, respectively, the work-based net cost of a step is given by E_step_ = (W^+^/e_1_) + (W^−^/e_2_). We use e_1_ = 0.326 and e_2_ = 1.085. These values are obtained by equally scaling the human muscle work efficiencies (25% and 125% for positive and negative work, respectively, Ruina et al 2003) such that the resulting minimum COT for walking at normal gravity and V = 1 ms^−1^ match the reported metabolic COT for the same conditions in Fig. 1 (simply to facilitate comparison of the model predictions with observations from human subjects moving under the same conditions).

Model predictions are obtained for impulsive actuation of all actuators. This is the optimal actuation profile when minimizing a work-based energetic cost with no bounds on the actuator forces/torques (Hasaneini et al., 2013, Srinivasan and Ruina, 2006). If upper bounds are imposed on actuator forces/torques, their rate of change is bounded or penalized, or forces/torques are expressed by a finite number of discrete points, the zero-duration infinite-magnitude impulses are replaced with more realistic burst forces/torques. However, in any of these cases, the general shape of the outputs (cost and step length curves for the study here) are preserved, although the resulting slopes will be slightly different. For example, similar trends are observed in preliminary results (Bertram and Hasaneini, 2013, Hasaneini, 2014), where a finite discretization scheme is used to express the actuator forces/torques.

At each gravity level, we find optimal walking at V = 1, 1.3, 1.6 m/s, and optimal running at V = 3 m/s by minimizing the COT (as defined above). For each walking gait, we consider that all actuators are non-regenerative (energy cannot be stored and recovered later in the gait cycle). For the running gaits, however, we calculate the push-off work using two different methods: (i) for a non-regenerative stance-leg actuator, and (ii) by assuming that 27% of the positive push-off work in each step is performed at no cost—as if it is recovered from the elastic energy stored in the muscles during the previous touch-down. We use 27% simply to match the scaling of the COT of the resulting optimal running at normal gravity and V = 3 m/s with that of humans for similar conditions (see Discussion). Although no rigorous analyses were used to determine the 27% elastic regeneration, it nonetheless provides a simple basis for examining the potential interplay between elasticity and gravity level. Not surprisingly, these two methods give two different optimal running gaits with different COT’s.

### Empirical measurements

#### Gravity reduction apparatus

We used a novel apparatus to simulate partial CoM gravity reduction. Here we explain the system and the logic behind its construction.

Following several other gravity simulation studies (e.g. Farley and McMahon, 1992, Donelan and Kram, 2000, Kram et al., 1997, Griffin et al., 1999) we simulate gravity changes with an approximately constant vertical force applied to a harness attached around the lower trunk (pelvis). In order to minimize use of lab space, and to reduce variation of force due to spring stiffness and material hysteresis, we have slightly modified the commonly utilized design. Instead of using extensive rubber springs with a small spring constant (employed to minimize force variation with length change), we use a mechanism that converts the linear spring into a (nonlinear) constant-force spring.

The general concept, discussed in Routh (2013) and reviewed in relation to the present mechanism by Herder (2001), is based on ‘a-static’ equilibrium, where equilibrium is maintained in a continuum of configurations. The zero rest-length spring, which is the a-static mechanism we use, was first invented by George Carwardine in 1931 for his Anglepoise lamp (Carardine, 1931, 1932), the design crudely copied in the two-parallelogram mechanism of modern student lamps. In seismology literature, the same mechanism is attributed to LaCoste (1934, 1935) and is used in the famous Press-Ewing long period seismometer (Press et al., 1958).

The apparatus is schematically described in Fig. 3. A hook pulls upward on a harness attached to the lower trunk of the subject. Straps connecting the harness to the hook are held free of the body by lightweight struts (tubular PVC plastic with large voids for weight reduction). The hook-harness assembly is suspended by a cable-pulley system, which itself is attached to a small trolley running on low friction wheels along one flange of an overhead I-beam. The moving cart mechanism reduces the horizontal force applied to the subject by the apparatus (compared to a fixed-hanging-point system), and allows for a more convenient horizontal displacement as the subject moves on the treadmill. This provides enough freedom to the subject for step length adjustment as she/he prefers.

**Figure 3:**
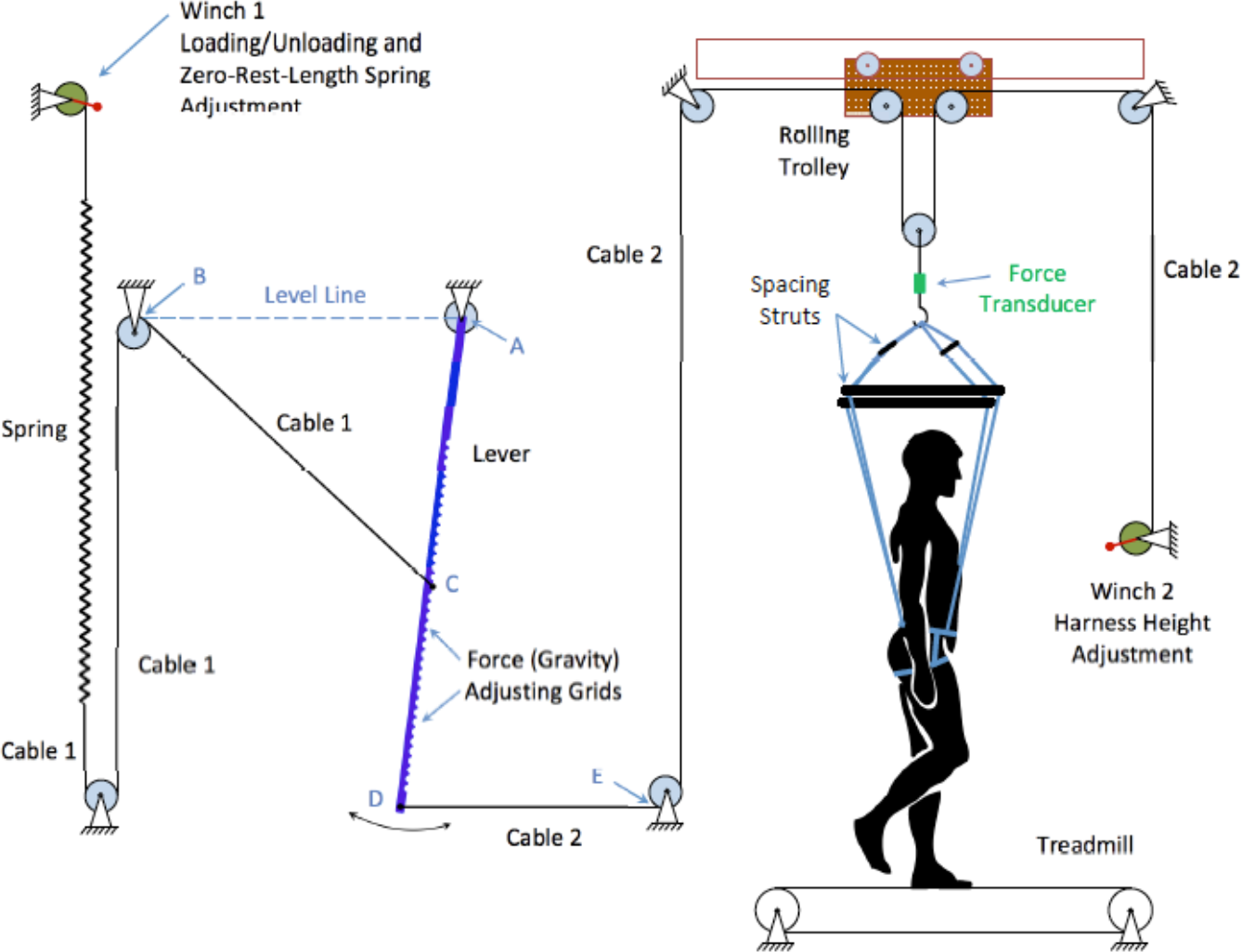
Reduced gravity apparatus based on zero-rest-length spring. A nearly-constant upward force is applied at the subject’s hip via a harness. This force magnitude can be adjusted by moving the cable-lever attachment point (point C) along the lever. The length of Cable1 and the Winch1 should be adjusted so that when the spring is at rest (Cable1 is not tensioned), the end of Cable1 attached at C will be at point B. Ideally, line DE should also be parallel to the level line AB. However, during the experiment, the subject’s hip/mass will have some vertical excursion, causing point D to move along the lever arc. Thus, to obtain the best performance, the dimensions of the system should be large enough to minimize the rotation of line DE. Also the lever moment of inertia about point A, the spring mass, and the friction of pulleys should be minimized to reduce variations of upward force during the gait. A rolling trolley is used to reduce the horizontal force applied to the subject and allow for minor horizontal displacement during the gait cycle. A force transducer is used to monitor and verify the upward force applied to the subject. Note that different elements of the figure are not to scale.

The functional mechanism consists of two separate cables, Cable1 and Cable2, connected to the pivoted rod AD (Fig. 3). Cable1 runs over multiple pulleys and connects the spring to the rod at point C, which can be repositioned to any of the slotted locations along the rod. Cable2 runs between Winch2 and the lower end of the rod, point D, via multiple pulleys, and is used to suspend the harness. During operation, the spring and subject’s weight hold both cables taught. The length of Cable1 is adjusted so that the spring is at its rest length if the end of Cable1 is detached from C and moved to B while Cable1 is still not slack (necessary to produce the zero rest-length spring function). Once the length of Cable1 is adjusted, Winch1 can be used to load/unload the system. The cable length is a critical factor in the function of the system (see below) so its length is carefully determined prior to use of the system. Due to the high loading levels involved (sometimes on the order of 75% body weight), Winch1 is conveniently used to reposition the spring-cable configuration while the subject remains in the harness.

Whereas the tension in Cable1 is proportional to the spring’s deflection, the tension in Cable2 is essentially constant. Neglecting the pulley and bearing friction, rod deformation, and inertial effects, the following rationale is presented. Denote the tension in Cable1 and Cable2 by T_1_ and T_2_, respectively. Because of the above-mentioned adjustment of the length of Cable1 (zero rest-length spring), we have T_1_ = k *ℓ*_BC_, where k is the spring constant and *ℓ*_BC_ is the length of the cable between B and C. Assume that the line DE remains parallel to the level line AB. The simplified geometry of the mechanism is shown in Fig. 4.

**Figure 4:**
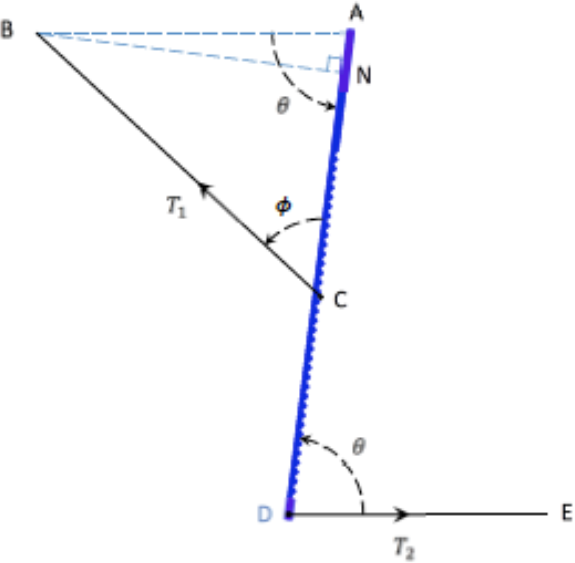
Simplified geometry of the moment balance features of the reduced gravity simulator. Line BN is normal to the lever. Ideally lines DE and AB are parallel.

Balancing moments about point A, we have:

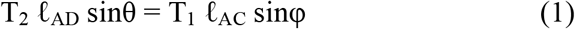

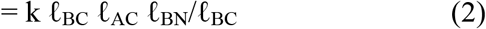

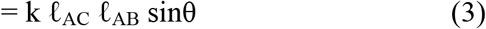

Therefore

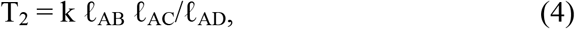
which does not depend on the lever angle θ and, subsequently, the vertical displacement of the subject. Also according to the above equation, the cable tension T_2_ can be adjusted by repositioning the point C along the lever (by adjusting *ℓ*_AC_).

In practice, as the harness moves up and down with the subject’s body position, the lever tip sweeps a small arc and the line DE slightly deviates from its ideal orientation (parallel to AB). This introduces some fluctuations in T_2_ within each step. Thus, to obtain the best performance, the dimensions of the system (lever length and the line DE) should be large enough, and the lever should be operated near vertical so that the rotation of line DE is minimized. Also the lever moment of inertia about point A, the spring mass, and the friction of pulleys should be as small as possible to minimize the variations of the upward force during the operation. In the Appendix we have calculated the cable tension T_2_ when DE is not necessarily parallel to AB.

### Subjects and study protocol

Subjects had no prior experience with reduced-gravity locomotion (note also that observations of humans in reduced gravity did not play a role in the construction of the model). All subjects were recreationally fit with no musculoskeletal or physiological impairments. Each subject provided informed consent approved by the University of Calgary Conjoint Health Research Ethics Board (CHREB) prior to participation.

### Study 1: Metabolic cost of normal and reduced gravity walking and running

Eight subjects participated in this portion of the study; 3 female, 5 male, age range 19 – 32 years (mean ± standard deviation; 24.1 ± 4.8). Height, leg length and body mass are listed in Table 1.

**Table 1.**
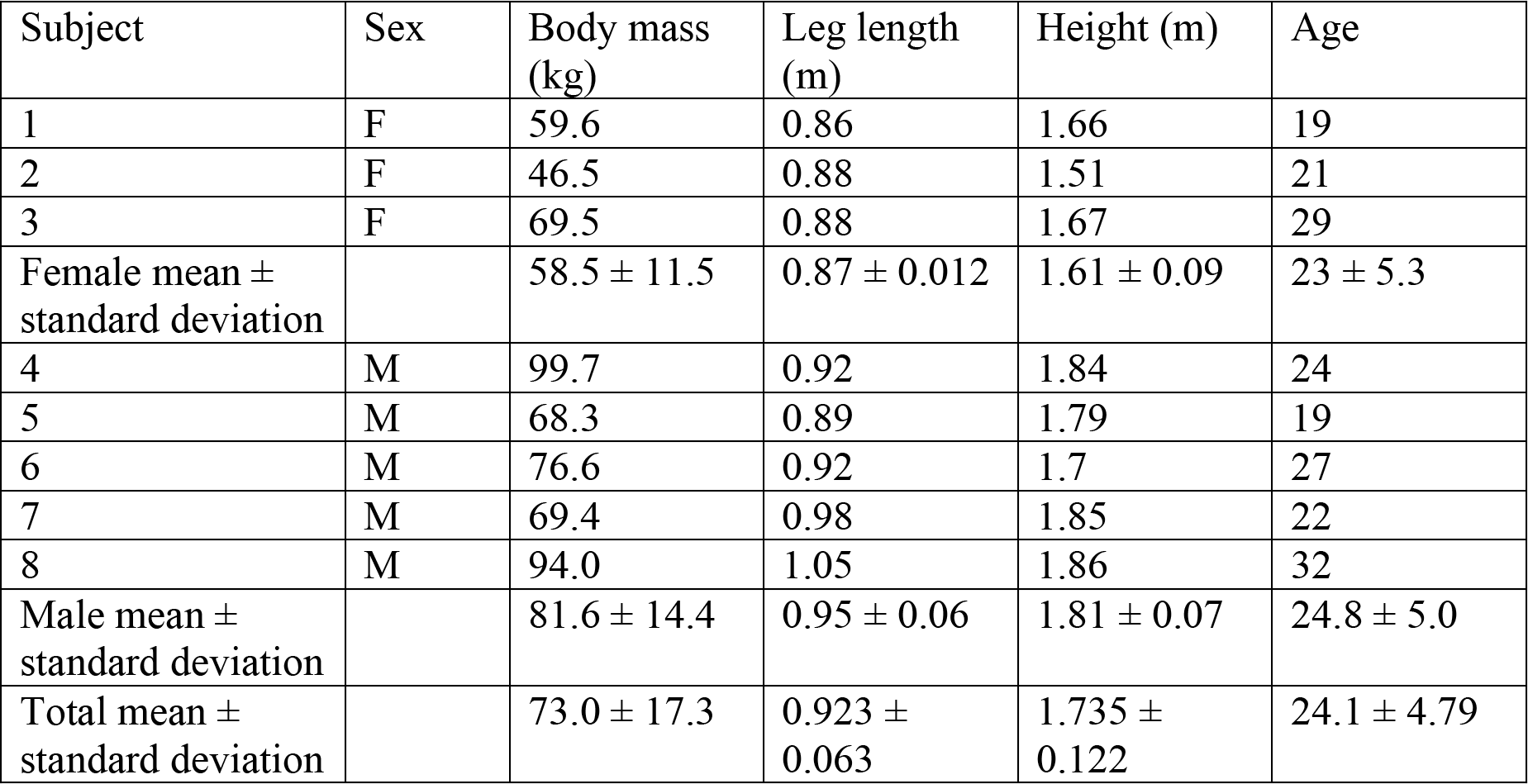
Characteristics of the subjects studied to evaluate the optimization model prediction of simulated gravity reduction on metabolic cost of locomotion.

| Subject | Sex | Body mass (kg) | Leg length (m) | Height (m) | Age |
| --- | --- | --- | --- | --- | --- |
| 1 | F | 59.6 | 0.86 | 1.66 | 19 |
| 2 | F | 46.5 | 0.88 | 1.51 | 21 |
| 3 | F | 69.5 | 0.88 | 1.67 | 29 |
| Female mean ± standard deviation | | $58.5 \pm 11.5$ | $0.87 \pm 0.012$ | $1.61 \pm 0.09$ | $23 \pm 5.3$ |
| 4 | M | 99.7 | 0.92 | 1.84 | 24 |
| 5 | M | 68.3 | 0.89 | 1.79 | 19 |
| 6 | M | 76.6 | 0.92 | 1.7 | 27 |
| 7 | M | 69.4 | 0.98 | 1.85 | 22 |
| 8 | M | 94.0 | 1.05 | 1.86 | 32 |
| Male mean ± standard deviation | | $81.6 \pm 14.4$ | $0.95 \pm 0.06$ | $1.81 \pm 0.07$ | $24.8 \pm 5.0$ |
| Total mean ± standard deviation | | $73.0 \pm 17.3$ | $0.923 \pm 0.063$ | $1.735 \pm 0.122$ | $24.1 \pm 4.79$ |

Oxygen consumption and CO_2_ elimination rates (STP) were determined using a commercial metabolic analysis system (TrueMax 2400, ParvoMedics, Salt Lake City, UT, USA). Measurements were considered acceptable if the respiratory exchange ratio (RER) remained below 1.0 for the final two minutes of each five to six minute trial.

For metabolic cost analysis two gravity levels were compared: normal (100%) and approximately 50% of normal (reduced gravity is approximate because the force-adjustment grid on the apparatus’ lever [see Fig. 3] has a resolution of about 15 N [magnitude of upward force]). Speeds were calculated based on the body size of the subject according to the square-root of the Froude number, Fr^0.5^ = (v^2^/(g_n_ L))^0.5^ where v is the treadmill velocity, g_n_ is normal gravitational acceleration (9.81 ms^−2^), and L is the subject’s leg length (ground surface to greater trochanter while standing upright wearing the sports shoes worn during the trials). At each gravity level subjects walked at three speeds and ran at three speeds. For a subject of leg length 0.90 m normal gravity walking speeds would be 1.0, 1.3 and 1.6 m/s (3.6, 4.7 and 5.8 km/hr) and running speeds would be 1.5, 2.2 and 3.0 m/s (5.4, 7.9 and 10.8 km/hr). In reduced gravity (≈ 50%) a subject with a 0.90 m leg length would have walked at 0.7, 1.0 and 1.3 m/s (2.5, 3.6 and 4.7 km/hr) and run at 1.0, 2.0 and 3.0 m/s (3.6, 7.2 and 10.8 km/hr). Normal gravity trials were conducted as a unit while the subject wore the harness, but the harness was not connected to the gravity simulation apparatus (while in an unloaded condition the loading straps and spacer bars moved unpredictably and interfered with the normal movement of the subject – this was not the case when the apparatus was under tension when simulating reduced gravity). The order of gait and speed were randomized for both normal and reduced gravity conditions. Prior to all trials, oxygen consumption and CO_2_ elimination rates (STP) were measured for each subject performing quiet standing for approximately a ten-minute duration.

### Study 2: Step length change with reduced gravity for walking and running

Sixteen subjects participated in this portion of the study; 6 female and 10 male, age range 19 – 36 years (mean ± standard deviation; 24.4 ± 5.3). The leg length (mean ± standard deviation) of the subject pool was *ℓ*_leg_ = 91.36 ± 4.7 cm, with body mass m = 67.55 ± 8.75 kg.

Each subject walked at three speeds (V_1_, V_2_, and V_3_) and ran at one speed (V_4_). Size differences among the subjects were accounted for as above, which allowed dynamically equivalent comparisons between individuals. In this portion of the study for
a subject with leg length *ℓ*_leg_ = 0.9 m, the speeds used would be V_1_ = 1 ms^−1^, V_2_ = 1.3 ms^−1^, V_3_ = 1.6 ms^−1^, and V_4_ = 3 ms^−1^.

At each speed the experiment was repeated for four different gravity levels. For 13 subjects these gravity levels were: normal gravity and approximately 80%, 60%, and 40% of normal. For the remaining three subjects the four gravity levels were: normal gravity and approximately 75%, 50% and 25% of normal. The latter set was used to extend the range and explore more extreme gravity reduction. We did not try this set for all subjects as we observed that most subjects did not feel comfortable in the harness at very low gravity levels (simulated low gravity requires large upward force applied to the harness that tends to destabilize some individuals). Actual simulated gravity levels were directly measured by the force transducer between the loading cable and the harness. Due to the limited resolution of the gravity-level adjustment grid on the lever, these varied slightly from the target values. The directly measured gravity level was used in all further analyses. Prior to data collection, subjects were given an opportunity to familiarize themselves with the system by applying an arbitrary reduced gravity simulation level different from that of the experimental trials.

For each subject the order of the twelve trials (three speeds at four gravity levels) was randomized. Once the harness load and treadmill speed were set for each trial, the subject was given time to become comfortable with the conditions presented. The subject self-determined this duration and individuals usually signaled their readiness for the trial within 20-40 seconds. During this period the treadmill speed and the contact patterns from foot sensors were visually monitored on a display to ensure that a steady state was achieved before data recording was initiated. Data collection continued for one minute in order to acquire adequate stride cycles to characterize gait under the applied conditions.

### Data acquisition

Upward force applied by the harness was monitored using a force transducer constructed by mounting strain gauges (Micro-Measurements, Raleigh, NC, CEA-06-125UW-350) in a half-bridge configuration on the tension and compression sides of a C-shape steel hook. This force-sensing hook was positioned as the connection between the tensioned cable and the harness (see Fig. 3). Following completion of all trials for a given subject, the force transducer (directly monitoring applied gravity level) was calibrated (scale factor and bias were calculated) using known weights.

To verify performance of the treadmill’s speed controller, and also to acquire more accurate speed information (e.g. within-step belt speed fluctuations caused by loading the treadmill motor with the subject’s body weight during foot contact), an optical encoder (Quantum Devices, Inc. Barneveld WI, QR12-24-0-ABLBA) and a custom-designed slope-estimator virtual instrument block in LabView (National Instruments, Austin, TX) were used to monitor the belt speed during the experiment.

For study 1, respiratory gas exchange was monitored continuously over the duration of each trial (5-6 minutes) to determine metabolic rate. The first three minutes allowed respiratory exchange to reach equilibrium under the imposed activity. This was assessed through visual inspection of the running average. If the operator was not satisfied that equilibrium had been reached, an additional minute was added to the trial. Energetic cost was calculated from the mean of the final two minutes providing the RER level was acceptable (RER < 1.0). Subjects were given an opportunity to rest between trials (gravity harness unloaded and metabolic mask removed).

Also for study 1, an inertial sensor (Xsens MTw, Xsens Technologies, Enschede, The Netherlands) was attached to each ankle. The acceleration data measured by these sensors were used to calculate the stride frequency in each trial. To measure step period in study 2, force sensitive resistors (Sparkfun Electronics, SEN-09375) were mounted directly to the skin surface of the heel and toe of each foot, where heel-strike and toe-off times could be detected separately (Snaterse et al., 2011). Direct measurement of foot contact (foot sensors) and belt speed (belt encoder) ensured accurate determination of step length variations, the measurement critical to the study 2 analysis.

A second optical encoder (QR12-2000-0-ABLBA) was installed on the rotational axis of the pivot rod of the gravity simulator (Point A, Fig. 3). Once calibrated, this provided a quantification of the vertical position of the harness since there is coupled motion between the pivot rod cable and the harness. This information was used to verify the correct functioning of the apparatus.

Analog (strain gauge and foot sensor) information was transferred to the signal conditioning circuit of a strain conditioning amplifier (National Instruments, Austin, TX,SCXI-1000 amp with SCXI-1520 8-channel universal strain gauge module connected through an SCXI-1314 terminal block) via flexible cables secured on the subject’s legs and running from the harness belt. The encoder’s digital output was fed through the SCXI system. The SCXI’s output was digitized (NI-USB-6251 mass termination) and acquired to a custom virtual instrument using LabView (National Instruments, Austin, TX).

### Data analysis

In the metabolic cost portion of the study (study 1) VO_2_ consumption rate of quiet standing was subtracted from the measurements, converted to units of Watts (assuming 20.1 Joules of energy released for each ml of O_2_ consumed) and then divided by the mass of the individual. This metabolic cost rate was then normalized as a metabolic cost of travel by converting the consumption rate to energy per distance traveled (metabolic rate divided by the average belt speed measured using the optical encoder over the corresponding trial period).

For the step length portion of the study (study 2) heel-strike and toe-off were detected from the voltage record of the foot-mounted sensors using a specified threshold signal level (transients were sharp and well defined for both the loading and unloading phases). Step times for each trial were generated from the time-differences between consecutive heel-strike or toe-off events for each leg (for each subject, speed and gravity level). The corresponding step period was then calculated by averaging the series of step times. Step length was obtained by multiplying the calculated step period and the average belt speed measured using the optical encoder over the corresponding trial period.

## Results

### Model predictions

The COT of energy-optimal gaits predicted by the model at different speeds and CoM-reduced gravity levels (i.e. partial reduced gravity) are shown in Fig. 5. The model predicts that gravity reduction has a greater influence on the energetics of running than walking. For all walking speeds measured, COT decreases slightly as gravity is reduced. For example, when comparing effective gravity conditions of normal and 20% of normal, the model’s COT for walking (V = 1 m/s) is reduced by 44%. The COT for running decreases by nearly twice as much (approximately 80%) over the same gravity range.

**Figure 5:**
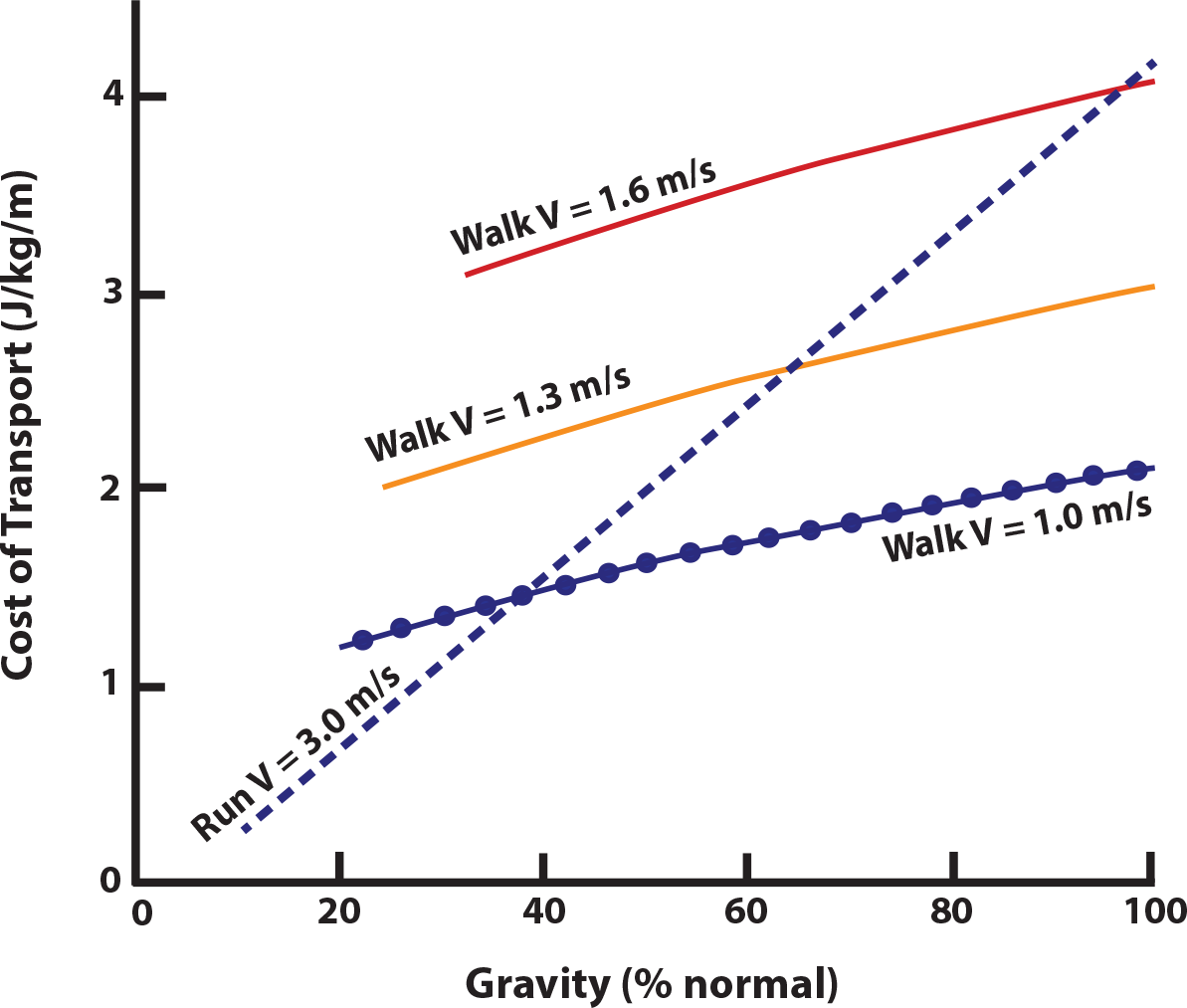
Cost of transport of energy-optimal gaits under partial reduced gravity conditions. Given the speed and gravity level, optimal gaits were obtained by minimizing the work-based energetic cost per unit distance traveled and unit body mass of the model shown in Fig. 2. Only the effective gravity on the center of mass is altered. Speeds correspond to those used for empirical measurements. No curve fitting is used in this figure (lines connect raw data points). Gait optimization predicts that the cost of running is more sensitive to gravity changes and this results in running having a lower cost of transport at low gravity levels, similar to that observed by Farley and McMahon (1992, Fig. 1).

Repeating the optimization for other running speeds results in overlapping cost relations because the COT for running is relatively insensitive to speed (see Fig. 6). The differential energetic response of walking and running to gravity leads the model to indicate that running should be more costly than walking at normal gravity but becomes less costly at lower gravity (a cost crossover occurs). Moreover, according to Fig. 5, as walking velocity decreases the crossover point occurs at lower gravity levels. This indirectly indicates that the preferred walk-run transition speed should decrease with reduced gravity, when speed is measured in absolute terms (Kram et al., 1997). The change in crossover point results from the differential energetic cost effects of gravity (running more sensitive) and speed (walking more sensitive).

**Figure 6:**
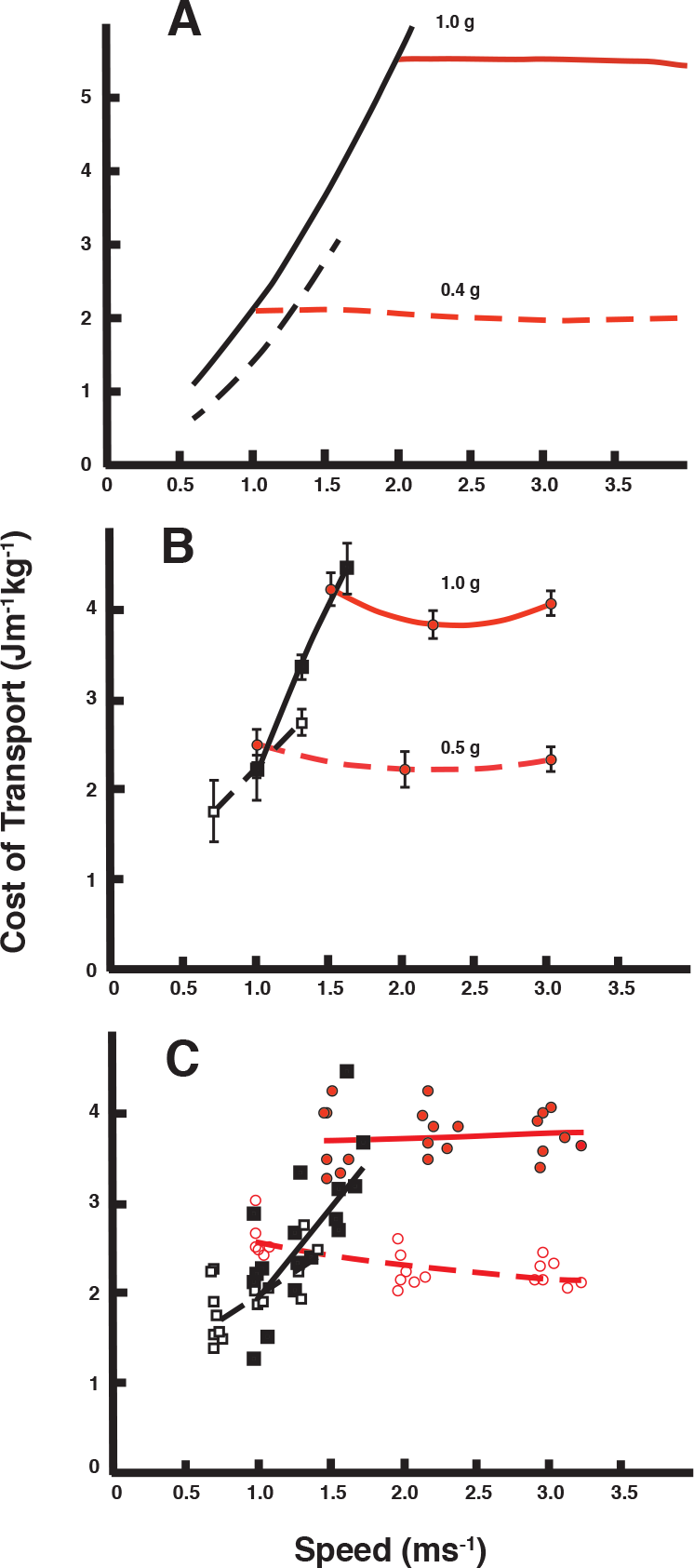
A. Model comparison of relationship between speed and cost of transport for two gravity levels (100% solid black line and 40% dashed black line) demonstrating a high sensitivity of walking transport cost to speed changes (lines have high slopes), but only modest effects for even a large change in gravity (lines are marginally displaced). This contrasts with running (1.0 g solid red line, 0.5 g dashed red line) where the cost of transport has low sensitivity to speed changes (low slope) but high sensitivity to gravity reduction (lines are well separated). B. Empirical measures parallel those of the model in a representative subject (points represent mean of three repetitions with error bars equaling 1 standard deviation; solid black squares, solid black line walk at 1.0 g, open squares, dashed black line walk at 0.5 g, red filled circles, red solid line run at 1.0 g, red open circles, red dashed line run at 0.5 g). C. Cost of transport for group of 8 subjects (see Table 1). Each point represents mean of three trials for each subject/condition. Symbols and lines as in B.

A slightly different view of these effects in walking and running energetics comes from plotting COT vs. speed for different gravity levels (Fig. 6A compares 40% g with normal gravity). The model predicts modest COT effects of gravity on walking but a strong influence for speed increases (a steep slope with 1.0 and 0.4 g plots residing adjacent to one another). This is contrary to running, in which gravity has a strong influence on COT while speed has little effect (low slope with substantially different COT magnitude at 1.0 and 0.4 g).

Optimization of the model for different speeds at reduced gravity levels also results in predictions of systematic step length changes (Fig. 7A). Although these effects are quite dramatic for running, walking exhibits much subtler trends. As such, they are plotted separately in Fig.7B with an appropriately scaled ordinate axis. As gravity is reduced, the model predicts that step lengths should increase for both energy optimal walking and running.

**Figure 7:**
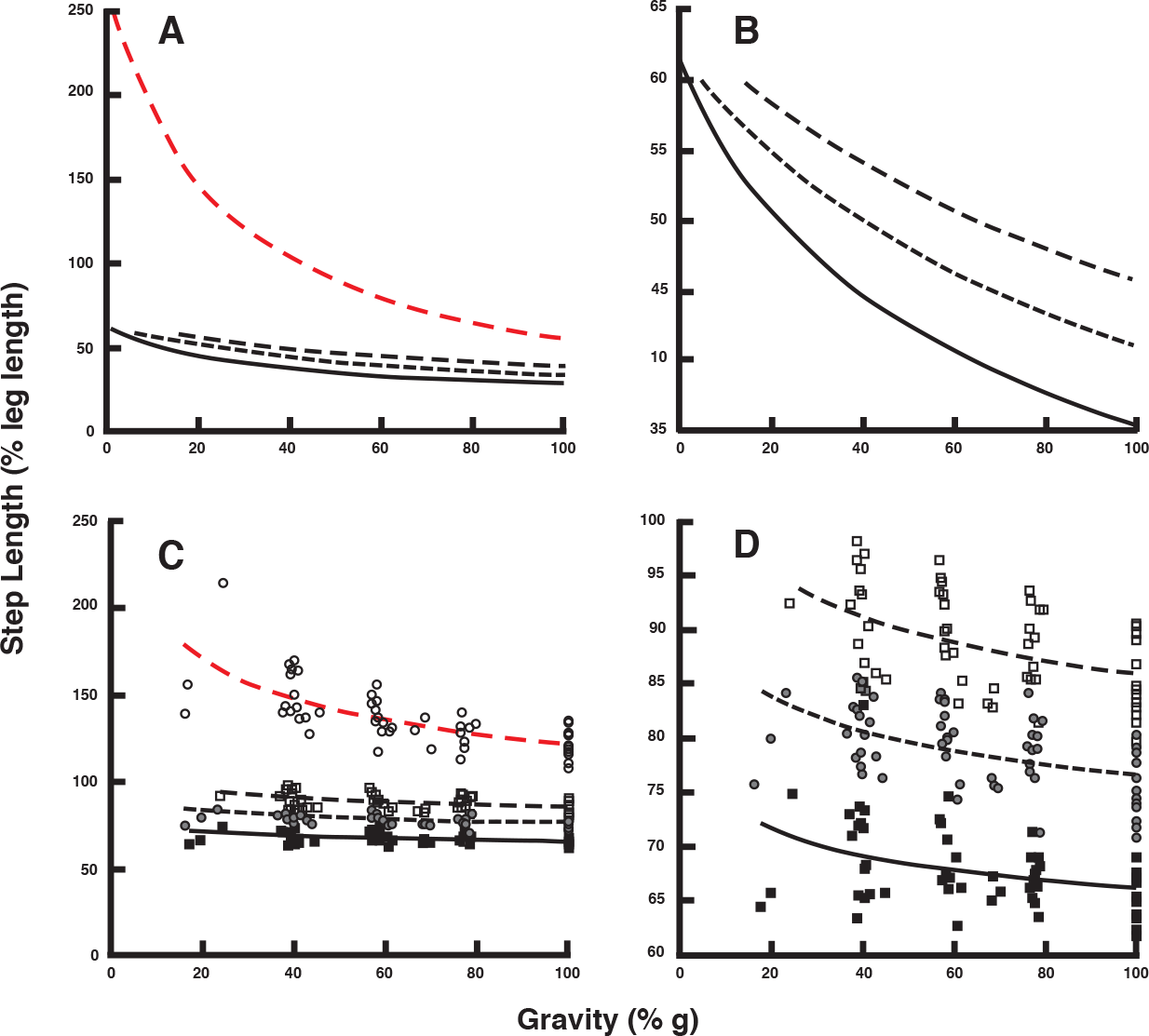
A. Comparison of the step length changes in walking and running as gravity level decreases from the optimization model. Red dashed line – run, and black lines – three walking speeds. B. The walking data from A with ordinate scales so that slopes are evident. C. Empirical measures from 16 subjects for conditions corresponding to A. D. Empirical measures for walking with ordinate scaled.

Cost of transport curves predicted by the model for faster walking do not reach to as low a gravity level as those for slower walking (Fig. 5). This occurs because the range of gravity levels for which walking is feasible decreases as speed increases. At faster speeds a larger centripetal force is required to keep the foot in contact with the ground while the center of mass moves on an almost circular path about the stance foot (the CoM moves as an inverted pendulum; Alexander, 1992, Usherwood, 2005). Normally this centripetal force should be provided by gravity, but when gravity is inadequate walking cannot be maintained and the biped is forced to switch to a running gait. In a similar manner, Fig. 7A and B depict the trend lines for all three walking speeds. They do not equally extend to very low gravity levels, but they do extend to gravity extremes that would not be comfortable for an individual. The lines are merely a prediction of the model, used to illustrate the trends only and do not indicate that we believe individuals can maintain normal walking to extremely low gravity levels at all speeds. Studies such as Cavagna et al. (2000) and Donelan and Kram (1997) show that the gravity range in which normal walking can be maintained decreases with speed.

The results reported above are for optimizations in which the effect of gravity was reduced on the CoM only, in order to be consistent with the experimental reduced gravity harness. We have verified that if gravity is reduced for the complete system in the model (full-body reduced gravity emulating true reduced gravity), the same energetics and step length trends are obtained with only slight changes in slope (note both gravity reduction methods give the same results at normal gravity, so differences at other gravity levels will emerge as a shift in modulation of slope).

### Empirical comparisons

Reliable empirical measures of COT for walking and running at constant speed over a range of simulated reduced gravity levels are available from Farley and McMahon (1992), so those measurements were not reproduced here (their results are shown in Fig. 1). We were interested in extending those data to generate a direct comparison of the model predictions for COT against walking and running velocity at two different gravity levels. Thus, empirical metabolic cost of transport at normal (1.0 g) and 50% normal (0.5 g) over a range of speeds for both walking and running are shown in Fig. 6B (single representative subject) and Fig. 6C (all eight subjects participating in this portion of the study). Farley and McMahon also presented similar results (their Fig. 2B).

The influence of gravity level on step length in walking and running is shown in Figure 7C (all 16 subjects participating in this portion of the study). Polynomial fits are plotted for the non-dimensionalized step length data of the entire subject pool. For better visualization of step length trends in walking, they are plotted separately in Fig. 7D. A polynomial regression is used for illustrative purposes only and is not meant to imply that step length changes with gravity according to this relation - only that it is non-linear and the polynomial is a convenient regression to demonstrate this. As these figures indicate, and in keeping with the predictions of the optimization model, the change in step length expected for walking is much less than that for running. Although step length increases in walking have shallow slopes, they all differ significantly from zero (p<0.0005; StatPlus, AnalystSoft, Inc.). The increasing step length trend is also valid for each subject individually, except for a few individual points at very low gravity levels.

## Discussion

This study compared the predictions of a minimally constrained optimization model of bipedal locomotion to human subject gait response for a range of speeds in an unusual functional environment – simulated reduced gravity. In so doing, the predictive capacity of the model is rigorously evaluated. In itself this is a valuable outcome because a predictive model that stands up to such scrutiny can serve as a valuable basis for more complex and realistic models that can be applied in a variety of areas, from understanding and interpreting pathological gait to suggesting novel robotic leg designs. The immediate objective of these studies, however, was to provide an explanation for why specific gait strategies are selected by the CNS. Developing a successful predictive model, even as simple as it was, allowed probing the model *post hoc* to determine the explicit consequences of each optimized movement strategy.

Model predictions for COT optimized walking and running were used to provide two specific evaluations of the model by comparing it with corresponding empirically measured features of walking and running in human subjects; the influence of speed and gravity on COT and accompanying step length changes. COT predictions for both running and walking compare favorably to the iconic results of Farley and McMahon (1992; compare Fig. 1 with Fig. 5) where the model anticipates that the COT in running is highly sensitive to gravity level, whereas those for walking change far less. This occurs in spite of the fact that the same model, actuators and cost function are used in both gaits. The difference is simply in the motion strategy expressed by the gait. Fig. 5 also indicates that differences in speed affect the COT of walking (note the different COT ranges for the three walking speeds shown). In contrast, the slope for running is quite steep, indicating a high sensitivity to gravity level. In Fig. 5 a single running plot is depicted because speed had such a small effect that model predictions overlapped substantially and would present as common set of lines that are difficult to visually distinguish. This is clearly evident in Fig. 6A where differences in gravity shift the COT of running while the slope over a range of speeds remains essentially horizontal.

The clear parallel between the model prediction and the empirical result (Fig. 6) is strong evidence that the two systems (model and human) are behaving in a similar manner. The inference, then, is that the motivation for that similarity is consistent between the two systems. Admittedly the specific magnitudes differ, but this is not surprising considering the simplicity of the model and the purposeful limitation in assumptions applied. However, the consistency in anticipated effects (trends) of two key functional factors, speed and gravity, each with converse effects on each gait, provides a compelling argument that COT optimization is an important determinant of the observed effects of speed and gravity level in human walking and running.

Model step length predictions also match the empirical results and these are consistent with several other reduced gravity studies where a variety of techniques to alter gravity level are used, such as parabolic flight path (full-body reduced gravity, Cavagna et al., 2000, De Witt et al., 2010) or a submersion method (Newman et al., 1994). However, our walking step length trends are in contrast to those observed by Donelan and Kram (2000), who used a spring-loaded apparatus to simulate reduced gravity conditions (although for running their observation is similar to ours). In their experiments subjects walked with slightly shorter step lengths as gravity level decreased. Because our results are consistent with those obtained with other reduced gravity simulation methods (Cavagna et al., 2000, De Witt et al., 2010, Newman et al., 1994) and the predictive model, we believe ours to be valid, but do not have an explanation for the contrary results of Donelan and Kram (2000).

Based on validation of the predictive capacity of the model, it is then possible to dissect the model *post hoc* to determine the physical basis (mechanisms responsible) for the COT effects and associated movement adjustments (such as step length).

Running: The optimization predicts that the supporting leg in running should make contact with the substrate in a near vertical position (Hasaneini et al., 2013). This minimizes dissipative (collision) loss because ground reaction force and CoM velocity vectors approach orthogonal. Dissipative loss is determined by the relationship between these these two vectors and that relationship will not be affected (much) by decrease in gravity, so the dissipative loss per step will be nearly constant. However, relief from downward acceleration during the non-contact phase of the running stride increases stride length, so an equivalent cost per contact with longer stride length means that running COT decreases directly with decrease in gravity level. At any given gravity level, though, change in speed involves an increase in dissipative loss (because COM velocity at contact increases) but this is offset by the concomitant increase in step length associated with faster running speeds (McMahon and Cheng, 1990); thus COT of running remains relatively insensitive to speed changes.

Walking: Speed increase in walking involves increase in stride length (Grieve and Gear, 1966). During the step-to-step transition from one support leg to the other the increase in COM velocity and change in leg contact geometry increases dissipative loss (Donelan et al., 2002), explaining the high energetic cost sensitivity of walking to speed. Walking cost remains relatively insensitive to decrease in gravity because these factors are unaffected by gravity – each stride progresses through contact of the new stance leg with the appropriate step length for the prescribed velocity, so mechanism and magnitude of loss is unaffected by gravity level. However, a decrease in gravity decreases the translation of kinetic and potential energy over the single limb stance portion of the step (inverted pendular motion). As a result, the decrease and subsequent increase in COM velocity of the pendular phase seen in normal gravity is reduced. The reduction of slowing over the stance phase that COM speed can be reduced slightly over the entire gait cycle even while translational speed is maintained. A reduction in COM speed would result in a decrease in dissipative loss at foot contact. However, the optimal strategy appears to be a subtle increase step length so that dissipative loss remains constant, but with slightly longer steps COT transport is decreased modestly.

In spite the model conceptually explaining the energetic cost trends associated with speed and gravity for walking and running, there are some aspects of the model’s optimization that do not match human gait. For example, the absolute value of the cost, the specific slope of the cost curves, and the gravity level where running becomes less costly than walking (the crossover point) are all quantitatively different between the optimization and empirical measurements. However, these details depend on many factors contributing to the complex human form and its physiology that are not included in the simple model. Although recognizing this, we take the similarity in the patterns involved, particularly with respect to the converse effects of both speed and gravity on the two gaits, as evidence that the objective function of the model represents a strong determinant of human locomotion strategy selection.

Farley and McMahon hypothesized that the differential response to gravity in walking and running is due to the different “energy-saving mechanisms associated with each of these gaits” (Farley and McMahon, 1992; p. 2711). They concluded that, “the link between the mechanics of locomotion and energetic cost is fundamentally different for running and for walking” (p. 2712). This hypothesis originates with the generally accepted perspective that the coordination patterns utilized in walking and running occur to maximize energy recovery within the stride cycle, e.g. Cavagna and Margaria (1966), Cavagna et al. (1977), Cavagna et al. (1964), Cavagna and Kaneko (1977), Cavagna et al. (1963) and He et al. (1991). According to this perspective, the best gait coordination strategy is the one that stores kinetic energy from one portion of the stride as some form of potential energy (gravitational for walking and strain potential energy for running) and then recovers it in another part of the cycle.

The energy recovery perspective is not supported by the optimization model results described here. Following the work reported in Srinivasan and Ruina (2006), we have previously shown that both walking and running emerge from work-cost minimization of a rigid model, with no direct or indirect reference to elasticity (Hasaneini et al., 2013). Here, we use the same model and energetic cost equation for both gaits, and still the optimization reasonably predicts the differential response trends for walking and running to both altered gravity and speed. This phenomenon manifests regardless of an absence of mechanisms capable of energy storage and recovery. Pendular exchange can be observed in walking and emerges as a consequence of the gait strategy used, not its objective. In fact, it is the reduction in pendular exchange that alters the dynamics of walking in reduced gravity that ultimately results in a modest reduction in COT at lower gravity levels.

Adding appropriate springs to the model for running gaits could decrease the cost of running and improve the estimation of such features as the energetic cost crossover gravity level. For instance, we have calculated the level of restitution (elastic recoil) required for the model cost of running in normal gravity to match the empirical measurements (Fig. 8), and the relationship generated is remarkably close to that found empirically over a broad range of reduced gravity conditions. This rough calculation suggests that approximately 30% (27% in our example) of the energy lost in stance contact would need to be returned for the model to match empirical observations. This shift in the cost along the relationship with gravity also makes the cross-over point between walking and running cost equivalent to that observed empirically in human subjects, i.e. as in Fig. 1. Note that this calculation does not represent true restitution included in the model, but is instead a simple *post hoc* calculation based on adding return to the cost relationship determined from the optimization model solution without restitution.

**Fig. 8.**
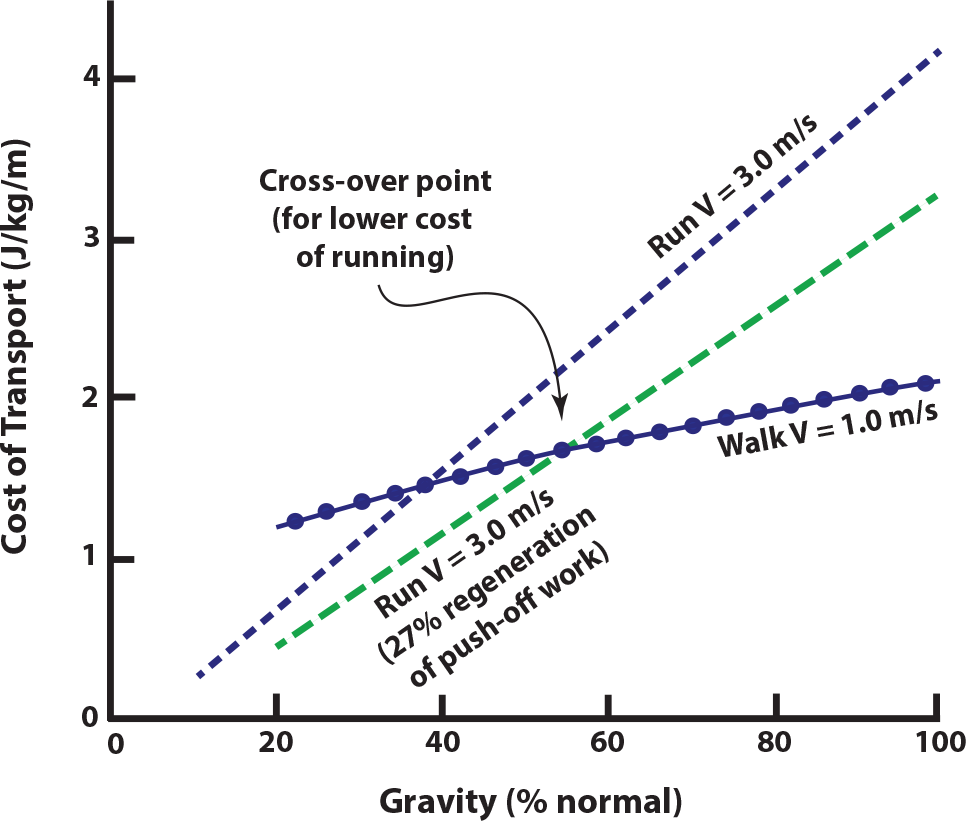
An estimate of the shift in running cost and change in cost crossover (where running at 3 m/s becomes less energetically costly than walking at 1 m/s) for the case of push-off work regeneration (27% of total push off work replaced in each step, green dashed line). This level of regeneration would shift the model running cost of transport curve and crossover point to match that observed by Farley and McMahon (1992).

As indicated by the above calculation, our disagreement with the recovery perspective as a key determinant of locomotion movement strategy does not imply that we question the occurrence or even advantage of energy storage and recovery in walking and running. Transfers between kinetic and various potential energy forms unequivocally occur in human and animal gaits through a variety of energy-saving mechanisms. It is also undeniable that these passive energy exchanges can reduce the energetic cost of locomotion by storing and reusing the energy at different phases of the gait. However, we argue these exchanges should be considered as descriptive characteristics of each gait, rather than regarded as a factor that determines the appropriate gait coordination strategy in each circumstance – just as foot contact overlap distinguishes human walking from running; overlap does occur, but it is merely a consequence of the different movement strategies that distinguish walking from running, not its cause.

We have shown that a minimally constrained model depending on simple but realistic dynamics of bipedal locomotion successfully predicts numerous aspects of human walking by optimizing for strategies that limit energetic loss and required leg work. The model parallels empirical observations of human locomotion, including the converse effects of speed and gravity on the energetics of walking and running. This occurs with the same model, actuators and cost function used in both gaits. Interestingly, the same simple model and optimization criterion also predicts many normally occurring features of human gait, including the optimality of walking at slow and running at fast speeds, the critically timed preemptive push-off in walking, landing on a near-vertical leg in running, the double maximum ground reaction force profile in walking and the single maxima for running, swing-leg retraction in walking and running, and the burst muscle activities at the start and end of various phases of motion in both gaits (Hasaneini et al., 2013). To this list can now be added the reduction of walk-run transition speed with decreases in gravity and the converse effects of speed and gravity on walking and running energetics. The ability of this simple model to successfully predict such encompassing aspects of human gait under both normal and unusual circumstances strongly suggests that the model optimization is determined by the same factors that influence human gait selection strategy. Through this we gain substantial insight into those features of gait and gait adaptation that appear valued by the CNS, the ultimate determiner of how gait is expressed.

1 In partial reduced gravity situations, the effective gravity level is reduced only on the body center of mass by applying a constant (ideally) upward force to the pelvis.

## References

Alexander, R. McN. (1992) A model of bipedal locomotion on compliant legs. Phil. Trans. R Soc. Lond. B 338, 189–198.

Alexander, R. McN. (2001). Design by numbers. Nature 412, 591.

Bekker, M.G. (1956). Theory of land locomotion. University of Michigan Press, Ann Arbor.

Bertram, J.E.A. (2005). Constrained optimization in human walking: cost minimization and gait plasticity. J. Exp. Biol. 208, 979–991.

Bertram, JEA, Ruina, A (2001) Multiple walking speed-frequency relations are predicted by constrained optimization J. Theor. Biol. 209, 445–453.

Bertram, J.E.A. and Hasaneini, S.J. (2013). Neglected losses and key costs: tracking the energetics of walking and running. J. Exp. Biol. 216, 933–938.

Biewener, A.A. (2006). Patterns of mechanical energy change in tetrapod gait: pendula, springs and work. J. Exp. Zool A Comp. Exp. Biol. 305(11), 899–911.

Borelli, G.A. (2012). On the movement of animals. Springer Science & Business Media.

Cavagna, G.A. and Margaria, R. (1966). Mechanics of walking. J. Appl. Physiol. 21(1), 271–278.

Cavagna, G.A., Heglund, N. and Taylor, C.R. (1977). Mechanical work in terrestral locomotion: Two basic mechanisms for minimizing energy expenditure. Am. J. Physiol. 233, 243–261.

Cavagna, G.A., Saibene, F. and Margaria, R. (1964). Mechanical work in running. J. Appl. Physiol. 19(2), 249–256.

Cavagna, G.A. and Kaneko, M. (1977). Mechanical work and efficiency in level walking and running. J. Physiol. 268, 467–481.

Cavagna, G.A., Saibene, F. and Margaria, R. (1963). External work in walking. J. Appl. Physiol. 18(1), 1–9.

Cavagna, G.A., Willems, P. and Heglund, N.C. (2000). The role of gravity in human walking: pendular energy exchange, external work and optimal speed. J. Physiol. 528(3), 657–668.

Cavagna, G.A., Heglund, N.C. and Taylor, C.R. (1977). Mechanical work in terrestrial locomotion: two basic mechanisms for minimizing energy expenditure. Am. J. Physiol. 233(5), R243–261.

Carwardine, G. (1932). Improvements in elastic equipoising mechanisms, UK Patent 404,615, (granted 4 January 1934).

Carwardine, G. (1931). Improvements in equipoising mechanism, UK Patent 379,680, (granted 22 August 1932).

De Witt, J.K., Perusek, G.P., Lewandowski, B.E., Gilkey, K.M., Savina, M.C., Samorezov, S. and Edwards, W.B. (2010). Locomotion in simulated and real microgravity: horizontal suspension vs. parabolic flight. Aviat. Space Environ. Med. 81(12),1092–1099.

Donelan, J.M. and Kram, R. (2000). Exploring dynamic similarity in human running using simulated reduced gravity. J. Exp. Biol. 203, 2405–2415.

Donelan, J.M. and Kram, R. (1997). The effect of reduced gravity on the kinematics of human walking: a test of the dynamic similarity hypothesis for locomotion. J. Exp. Biol. 200, 3193–3201.

Donelan, J.M, Kram, R. and Kuo, A.D. (2002) Mechanical work for step-to-step transitions is a major determinant of metabolic cost of human walking. J. Exp. Biol. 205, 3717–3727.

Farley, C. and McMahon, T.A. (1992). Energetics of walking and running: insights from simulated reduced-gravity experiments. J. Appl. Physiol. 73, 2709–2712.

Gill, P.E., Murray, W. and Saunders, M.A. (2005). SNOPT: An SQP algorithm for large-scale constrained optimization. SIAM Rev. 47, 99–131.

Grieve, D.W. and Gear, R.J. (1966) The relationship between length of stride, step frequency, time of swing and speed of walking for children and adults. Ergonomics 9(5): 379–399.

Griffin, T.M., Tolani, N.A. and Kram, R. (1999). Walking in simulated reduced gravity: mechanical energy fluctuations and exchange. J. Appl. Physiol. 86(1), 383–390.

Gutmann, A., Jacobi, B., Butcher, M.T. and Bertram, J.E.A. (2006). Constrained optimization in human running. J. Exp. Biol. 209, 622–632.

Hasaneini, S.J. Energy efficient bipedal locomotion. PhD thesis, University of Calgary, Calgary, Alberta, Canada, Jan 2014.

Hasaneini, SJ, Bertram, JEA, Macnab, CJB (2015) Energy optimal relative timing of stance-leg push-off and swing-leg retraction in walking. Robotica (doi:10.1017/S0263574715000764).

Hasaneini, S.J., Macnab, C.J.B., Bertram, J.E.A. and Leung, H. (2013). The dynamic optimization approach to locomotion dynamics: human-like gaits from a minimally-constrained biped model. Adv. Robot. 27(11), 845–859.

He, J., Kram, R. and McMahon, T.A. (1991). Mechanics of running under simulated low gravity, J. Appl. Physiol. 71(3), 863–870.

Herder, J.L. (2001). Energy-free Systems. Theory, conception and design of statically. PhD thesis, Technical University of Delft, Delft, Netherlands.

Kuo, A.D. (2001). A simple model of bipedal walking predicts the preferred speed-step length relationship. J. Biomech. Eng. 123(3), 264–269.

Kuo, A. D. (2002). Energetics of actively powered locomotion using the simplest walking model, J. Biomed. Eng. 124, 113–120.

Kram, R., Domingo, A. and Ferris, D.P. (1997). Effect of reduced gravity on the preferred walk-run transition speed. J. Exp. Biol. 200(4), 821–826.

LaCosteJr, L.J. (1934). A new type long period vertical seismograph. J. Appl. Physics 5(7), 178–180.

LaCoste, L.J. (1935). A simplification in the conditions for the zero-length-spring seismograph. Bull. Seismol. Soc. Am. 25(2), 176–179.

Lee C.R. and Farley, C.T. (1998). Determinants of the center of mass trajectory in human walking and running. J. Exp. Biol. 201(21), 2935–2944.

Lee, DV, Comanescu, TN, Butcher, MT, Bertram, JEA (2013) A comparative collision-based analysis of human gait. Proc. Roy. Soc. Lond. B 280(1771): (20131779).

Margaria, R, Cerretelli, P, Aghemo, P, Sassi, G (1963) Energy cost of running. J. Appl. Physiol. 18(2): 367–370.

McGeer, T. (1990a). Passive dynamic walking. Int. J. Robot. Res. 9, 68–82.

McGeer, T. (1990b). Passive dynamic running. Proc. Roy. Soc. Lond. B 240, 107–134.

McMahon, T.A. and Cheng, G.C. (1990) The mechanics of running: How does stiffness couple with speed? J. Biomech. 23(S1), 65–78.

Newman, D.J., Alexander, H.L. and Webbon, B.W. (1994). Energetics and mechanics for partial gravity locomotion. Aviat. Space Environ. Med. 65(9), 815–823.

Polet, D.T., Schroeder, R.T. and Bertram, J.E.A. (2017) Gravity takes the bounce out of running: work-based optimization explains flatter gait in reduced gravity. J. Exp. Biol. (submitted).

Press, F., Ewing, M. and Lehner, F. (1958). A long-period seismograph system. Eos, Trans. Am. Geophys. Un. 39(1), 106–108.

Pribe, C., Grossberg, S. and Cohen, M.A. (1997). Neural control of interlimb oscillations. II. Biped and quadruped gaits and bifurcations. Biol. Cybern. 77, 141–152.

Rashevsky, N. (1948). On the locomotion of mammals. Bull. Math. Biol. 10, 11–23.

Rashevsky, N. (1960). Mathematical Biophysics: Physico-mathematical foundations of biology. Form and locomotion in some quadrupeds, pp. 262–269. Vol. 2, 3^rd^ revised ed. Dover Pub, New York.

Romijn, J.A., Coyle, E.F., Hibbert, J. and Wolfe, R.R. (1992). Comparison of indirect calorimetry and a new breath ^13^C/^12^C ratio method during strenuous exercise. Am. J. Physiol. Endocrinol. Metab. 263, E64–E71.

Routh, E.J. (2013). A Treatise on Analytical Statics: With Numerous Examples, vol. 1. Cambridge University Press.

Ruina, A., Bertram, J.E.A. and Srinivasan, M. (2005). A collisional model of the energetic cost of support work qualitatively explains leg sequencing in walking and galloping, pseudo-elastic leg behavior in running and the walk-to-run transition. J. Theor. Biol. 14(2), 170–192.

Saibene, F. (1990). The mechanisms for minimizing energy expenditure in human locomotion. Eur. J. Clin. Nutr. 44(Suppl. 1), 65–71.

Selinger, J.C., O’Connor, S.M., Wong, J.D. and Donelan, J.M. (2015) Humans can continuously optimize energetic cost during walking. Curr. Biol. 25(18), 2452–2456.

Snaterse, M., Ton, R., Kuo, A.D. and Donelan, J.M. (2011). Distinct fast and slow processes contribute to the selection of preferred step frequency during human walking. J. Appl. Physiol. 110(6), 1682–1690.

Srinivasan, M. and Ruina, A. (2006). Computer optimization of a minimal biped model discovers walking and running. Nature 439(7072), 72–75.

Tucker, V.A. (1975). The energetic cost of moving about. Am. Sci. 63, 413–419.

Usherwood, J.R. (2005) Why not walk faster? Biol. Lett. 1(3), 338–341.

Winter, D. (2005). Biomechanics and motor control of human movement, pp. 60–64. John Wiley & Sons, Inc.

Wong, J.D. and Donelan, J.M. (2017) Principles of energetics and stability in human locomotion. SemanticScholar (https://www.semanticscholar.org/paper/Principles-of-energetics-and-stability-in-human-Wong-Donelan/ff1c1adbbd829ed781158c21fb7bdc17cbdcdf9a

